# GLARE: Genomic Language model for Automated Repetitive Element classification

**DOI:** 10.64898/2026.10.06.757139

**Authors:** Nicolas Haas, Tomáš Brůna

## Abstract

Reliable classification of transposable elements (TEs) is important for genome annotation and evolutionary analysis, yet remains challenging: TE families differ widely across taxa, reference databases label them inconsistently, and many TEs are represented only by degraded copies. We introduce GLARE (**G**enomic **L**anguage model for **A**utomated **R**epetitive **E**lement classification), a TE classifier built by fine-tuning a genomic language model on harmonized PanTEon and Repbase reference data to predict TE superfamily across 32 superfamilies in 11 orders. Among the genomic language models compared, Nucleotide Transformer v3 provided the strongest representation. Built on this backbone, GLARE exceeded all nine classifiers in the PanTEon benchmark, reaching 0.966 weighted superfamily F1 on its 30-superfamily task. GLARE also showed stronger generalization to clades absent from training, consistently exceeding state-of-the-art classifiers on holdouts spanning three taxonomic ranks: the order Poales, the phylum Arthropoda, and the kingdom Fungi. Training on both PanTEon and Repbase reduced the performance drop seen when a single-source model was applied to the other database. Under simulated sequence decay, GLARE was the most robust classifier tested, and it performed best on independently curated degraded libraries. These findings show that genomic language models fine-tuned on harmonized multi-source references can classify TEs robustly under the taxonomic, database, and sequence-quality shifts that matter in practice. GLARE is available at code.jgi.doe.gov/NHaas/glare.

## 1 Introduction

Transposable elements (TEs) are mobile DNA sequences that make up a large and highly variable share of eukaryotic genomes. New insertions create genetic variation, and accumulated copies provide substrates for recombination and drive sequence turnover [O’Donnell and Burns, 2010]. As a result, TEs are major contributors to differences in genome size and architecture. Furthermore, TE-derived sequences can be co-opted into regulatory programs [Bourque et al., 2018, Feschotte, 2008, Chuong et al., 2017]. These effects make TEs informative records as well as active drivers of genome evolution. However, their abundance and repetitive nature complicate genome assembly, gene annotation, and comparative analysis [Makałowski et al., 2019, Goerner-Potvin and Bourque, 2018]. For these reasons, accurate TE identification and classification are essential not only for investigating TE biology, but also for virtually any genome-scale analysis, such as reconstructing evolutionary history or making meaningful comparisons across genomes.

TEs are classified hierarchically according to their transposition mechanism, structural features, and evolutionary relationships. In the widely used Wicker system, TEs are categorized into two main classes: Class I retrotransposons mobilize via a copy-and-paste mechanism using an RNA intermediate, whereas Class II elements use DNA-based mechanisms. Classes are divided into orders, some of which are further divided into superfamilies [Wicker et al., 2007]. Repbase, the most widely used transposable element reference database, and other resources use related but not identical nomenclatures—their systems differ in how they accommodate non-autonomous, chimeric, and newly discovered elements [Bao et al., 2015, Piégu et al., 2015]. Even within one system, superfamily assignment is often hard in practice. Many copies are fragmented or degraded, and others are nested within or recombined with other elements [Makałowski et al., 2019, Loreto et al., 2024]. Comparisons between libraries or classifiers therefore need to state which labels they use and at what level.

TE classification typically relies on curated reference libraries. Repbase distributes consensus sequences of repetitive elements, whereas Dfam represents TE families as profile hidden Markov models [Bao et al., 2015, Storer et al., 2021]. Annotation pipelines such as RepeatModeler2 can discover TE families *de novo* [Flynn et al., 2020], but the resulting consensus sequences must still be assigned to classes, orders, and superfamilies, and many remain unclassified. Most pipelines label these sequences by homology to Repbase or Dfam; RepeatModeler2, for instance, does this with RepeatClassifier. Standalone classifiers also search for diagnostic features. REPCLASS combines homology and structural searches, and PASTEC uses protein-domain profiles together with features such as long terminal repeats (LTR) and terminal inverted repeats (TIR) [Feschotte et al., 2009, Hoede et al., 2014]. The rule- and homology-based approaches are interpretable, since each label can be traced to a specific match or feature. However, they cannot reliably classify elements that lack close relatives in reference libraries or whose diagnostic features have been lost through truncation or decay.

Machine learning offers an alternative route from sequence to hierarchical labels. TEclass is an early example, classifying sequences from their oligomer frequencies with support vector machines [Abrusán et al., 2009]. Later methods cover more levels of the TE hierarchy and increasingly rely on neural networks. Many still start from *k*-mer counts: ClassifyTE combines *k*-mer features in a stack of classifiers [Panta et al., 2021], and DeepTE feeds *k*-mer-derived inputs into hierarchical convolutional neural networks (CNNs) [Yan et al., 2020]. TERL, which also relies on CNNs, instead learns directly from one-hot-encoded sequences [da Cruz et al., 2020]. Inpactor2, developed for plant genomes, builds a classifier into an annotation pipeline that detects candidate LTR retrotransposons, discards partial elements, and assigns intact ones to superfamilies and lineages [Orozco-Arias et al., 2023]. Recent methods differ more widely in both features and architectures. NeuralTE combines *k*-mer patterns from the internal and terminal regions of each sequence with features representing target site duplications, terminal repeats, and protein domains [Hu et al., 2024]. CREATE integrates global *k*-mer distributions with terminal sequence profiles in a network of convolutional, recurrent, and attention layers [Qi et al., 2025]. Others borrow from language modeling. BERTE combines representations from a BERT encoder pretrained on natural language [Devlin et al., 2018] with *k*-mer frequencies [Chen et al., 2024], and TEclass2 applies a transformer to tokenized consensus sequences [Bickmann et al., 2025]. Terrier, a hierarchical convolutional model that predicts order and superfamily labels, demonstrated that deep learning classifiers can substantially reduce the proportion of unclassified elements compared to traditional rule- and homology-based pipelines [Turnbull et al., 2025].

Although these machine learning methods broaden TE classification beyond direct homology, they differ in training databases, taxonomic composition, label hierarchies, and data splits, so their reported performance cannot be compared directly. PanTEon addresses many of these challenges with a harmonized TE collection spanning multiple kingdoms and a modular framework for training and evaluating classifiers under shared procedures [Orozco-Arias et al., 2026]. Its latest version also evaluates classifiers on unseen species and degraded TEs, but neither the evolutionary distance between training and test sequences nor the extent of degradation is controlled.

Genomic language models (gLMs) offer a complementary way to learn sequence representations that can be adapted to TE-specific tasks. gLMs are first pretrained on large collections of DNA with self-supervised objectives, such as predicting masked nucleotides. Through this pretraining, gLMs develop a contextual representation of DNA that can be subsequently fine-tuned for specific tasks, potentially reducing the need for hand-crafted classification features [Dalla-Torre et al., 2024, Zhou et al., 2023]. However, pretraining alone does not guarantee better performance than task-specific supervised models, or better generalization beyond the training data [Benegas et al., 2025]. Model performance depends on design choices such as the pretraining data, tokenization, architecture, context length, strand handling, and fine-tuning strategy. Two of these matter particularly for TEs: many elements are longer than a model’s context window, and a TE’s identity should not change when it is read from the opposite strand. The backbones evaluated in this study handle these issues differently: Caduceus is built to be reverse-complement equivariant and to model long sequences, DNABERT-2 uses byte-pair encoding to represent nucleotide motifs of variable length, and Nucleotide Transformer v3 (NTv3) extends multispecies pretraining to long contexts [Schiff et al., 2024, Zhou et al., 2023, Boshar et al., 2025].

Here, we introduce GLARE (**G**enomic **L**anguage model for **A**utomated **R**epetitive **E**lement classification), which fine-tunes the NTv3 genomic language model for TE classification. GLARE predicts TE superfamilies and derives orders from them, so labels at the two levels are always consistent. We first compare GLARE with nine classifiers within the PanTEon framework, where all methods are trained on the same data and scored the same way. We then test its generalization to clades withheld from training at the order, phylum, and kingdom levels, and assess its performance on incomplete and degraded elements, its transfer across databases, and its sensitivity to reverse-complement orientation. Across these evaluations, GLARE retained the highest accuracy on withheld clades and on degraded sequences, and outperformed all nine classifiers evaluated in the PanTEon benchmark.

## 2 Materials and Methods

### 2.1 The GLARE classifier

#### 2.1.1 Training data and label space

The final GLARE classifier was trained on the PanTEon sequence collection benchmark-edition v1.6.1 [Orozco-Arias et al., 2026] and Repbase release 31.03 [Bao et al., 2015]. All annotations were translated into a common vocabulary using explicit, source-specific mapping tables (Supplementary Data 1–2). This Wicker-compatible vocabulary [Wicker et al., 2007] comprised 32 superfamilies nested within 11 orders. Each label keeps both levels of the hierarchy, as in LTR/GYPSY or TIR/TC1MARINER. Source categories without a Wicker-compatible superfamily were assigned to the OTHER label of their parent order, and records whose annotation did not resolve to a single order were removed. Satellites and structural RNAs were kept because GLARE is intended for annotating complete *de novo* repeat libraries, where these sequences are abundant and often mistaken for TEs. The full vocabulary and hierarchy are listed in Table 1. This 32-superfamily vocabulary applies to the released model only; the comparative experiments in Section 2.2 use PanTEon’s native vocabulary of 30 superfamilies in 8 orders.

**Table 1:** Final 11-order, 32-superfamily label space of the released GLARE model. Each superfamily appears under exactly one order; predicted order is derived deterministically from the predicted superfamily through this map.

| Order | Superfamily labels |
| --- | --- |
| CRYPTON | CRYPTON |
| DIRS | DIRS |
| HELITRON | HELITRON |
| LINE | CR1; I; L1; OTHER; R1; RTE |
| LTR | BELPAO; COPIA; ERV; GYPSY; OTHER |
| MAVERICK | MAVERICK |
| PLE | PLE |
| SATELLITE | SATELLITE |
| SINE | 7SL; OTHER; TRNA |
| STRUCTURAL_RNA | STRUCTURAL_RNA |
| TIR | ACADEM-1; CACTA; HAT; KOLOBOK; MERLIN; MULE; OTHER; P;<br>PIFHARBINGER; PIGGYBAC; TC1MARINER |

*Note.* Comparative experiments (Section 2.2) use PanTEon’s native 30-superfamily, 8-order vocabulary.

After this mapping, the input set contained 230,428 PanTEon and 117,778 Repbase records. Each source was independently split by stratified random sampling, with 90% used for training and validation and 10% held out for testing, giving external test sets of 23,043 PanTEon and 11,778 Repbase records. The two train–validation portions were then pooled and split again, with stratification, into 282,046 training and 31,339 validation records. All partitions were assigned per record rather than per sequence-similarity cluster, matching the PanTEon benchmark, whose fixed record assignments are needed for the comparison in Section 2.2.1. As a result, near-identical copies from large families can fall into different partitions, so within-database scores are best read as upper bounds. The clade-holdout, unseen-species, and degradation tests described below exclude whole species or perturb held-out sequences, and therefore provide a stricter test of generalization.

#### 2.1.2 Architecture

GLARE (Figure 1) is built on the pretrained `InstaDeepAI/NTv3_650M_pre` Nucleotide Transformer v3 (NTv3) backbone [Boshar et al., 2025], chosen in the backbone comparison described in Section 2.3. All backbone parameters were fine-tuned. Token-level hidden states are reduced by masked mean pooling over non-padding positions. The pooled vector is passed to a classification head with two blocks. Each block consists of a linear projection, layer normalization, GELU activation, and dropout; the first reduces the backbone hidden size of 1,536 to 512 units, and the second reduces it to 256. A final linear layer outputs one logit per superfamily. The model was trained on superfamily labels only (Section 2.1.3), and orders are assigned from the predicted superfamily using the fixed superfamily-to-order map (Table 1).

**Figure 1.**
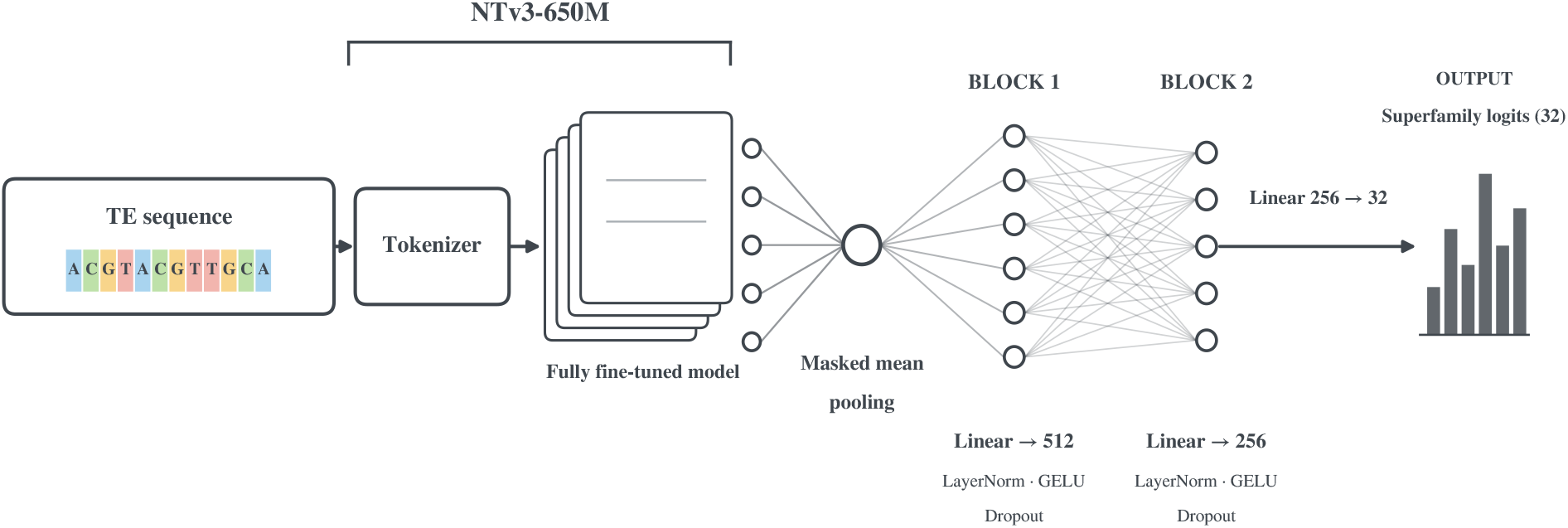
GLARE architecture. A TE sequence is tokenized with the NTv3 tokenizer and encoded by the fully fine-tuned NTv3-650M backbone. Token-level hidden states are mean-pooled over non-padding positions and passed through two classification blocks (1,536 → 512 and 512 → 256), each consisting of a linear projection, layer normalization, GELU activation, and dropout. A final 256 → 32 linear projection produces the superfamily logits.

#### 2.1.3 Training

Sequences were presented at a maximum context of 4,096 tokens. Longer sequences were represented by a uniformly sampled contiguous crop, and shorter sequences were padded by the tokenizer. Sampling one crop per example per epoch limits the cost of each step and serves as an implicit augmentation; at inference, all overlapping windows are used (Section 2.1.4). Reverse-complement augmentation replaced each sampled training example with its reverse complement with probability 0.5, leaving labels unchanged.

The classifier was optimized for superfamily prediction using weighted cross-entropy with label smoothing, where the class weights were square-root inverse class frequencies normalized to mean one among classes represented in the training set. Optimization used AdamW with separate parameter groups for the backbone and classification head, a cosine learning-rate schedule, and linear warmup over the first 5% of optimizer steps. Training used BF16 CUDA autocasting, gradient accumulation over four minibatches, and gradient clipping at Euclidean norm 1.0. Training was limited to ten epochs, and the checkpoint with the highest validation weighted superfamily F1 was retained.

After checkpoint selection, a single temperature *T* was fitted to the validation superfamily logits by minimizing multiclass negative log-likelihood [Guo et al., 2017]. The confidence threshold described below (Section 2.1.4) is applied to the resulting calibrated probabilities.

#### 2.1.4 Inference

Sequences no longer than the context length *L* are classified as a single window. Longer sequences are split into overlapping windows of length *L*, starting at the first nucleotide and advancing by max{1, ⌊*L*(1 ™ *o*)⌋}, where *o* is the overlap fraction. The final window may be shorter than *L*. Each window is tokenized separately, and the logits of all windows from a record are averaged. We used *L* = 4,096 and *o* = 0.5, giving a stride of 2,048 nt.

The averaged logits *z*_*k*_ are then scaled by the fitted temperature (*T* = 0.721) and converted to probabilities:

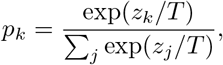

GLARE reports the superfamily with the highest probability, uses that probability as its confidence, and assigns the order from the superfamily-to-order map. By default, predictions with confidence below 0.7 are labeled `UNKNOWN/UNKNOWN.`

Temperature scaling does not change which superfamily scores highest, so it has no effect on the F1 values reported here; the threshold, which can change the reported label, was disabled for all comparative evaluations (Section 2.2).

#### 2.1.5 Hyperparameter optimization

Hyperparameters were selected with an Optuna study [Akiba et al., 2019] using the Tree-structured Parzen Estimator sampler [Watanabe, 2023] over 40 trials. Each trial was trained for at most ten epochs and was ranked by its highest validation weighted superfamily F1. The search covered the backbone and head learning rates, weight decay, batch size, label smoothing, class-weighting scheme, context length, validation-window overlap, hidden dimension, and head dropout. Supplementary Table S1 reports the search domain and the selected value for each optimized dimension. The validation records used for hyperparameter selection were disjoint from both the final test partition and the PanTEon benchmark test sets.

### 2.2 Accuracy evaluation

#### 2.2.1 Comparison with existing TE classifiers on PanTEon benchmark

To compare GLARE with existing TE classifiers, we evaluated it on the PanTEon benchmark, which retrains each classifier on the same training data [Orozco-Arias et al., 2026]. The benchmark includes nine classifier architectures evaluated on a 30-superfamily multiclass task, and their reported F1 values serve as the comparison. PanTEon trains classifiers on a cross-kingdom database with 80/10/10 training/validation/test partitions, and reports test-set class-support-weighted multiclass F1. GLARE was evaluated under the PanTEon conditions, using the record assignments distributed with the benchmark. The GLARE values reported for this benchmark therefore derive from the same held-out sequences, scored with the same weighted multiclass F1 implementation, and from a model trained on an identical training set.

We additionally evaluated GLARE on the unseen-species evaluation sets curated by the PanTEon authors from Dfam v4.0 [Storer et al., 2021, Orozco-Arias et al., 2026]. Briefly, this curation involved the removal of all sequences belonging to species represented in the PanTEon training set from the complete curated Dfam collection. Non-TE repeats, including satellites and simple repeats, were excluded. For LTR retrotransposons, matching LTR and internal regions were merged by sequence identifier whenever possible, yielding 2,309 TE sequences. PanTEon’s workflow separated these records into 867 complete elements satisfying the expected structural requirements and 1,442 incomplete or degraded elements.

For the benchmarking experiments that follow, we selected NeuralTE and Terrier, the two highest-ranked models across all of PanTEon’s benchmarks.

#### 2.2.2 Generalization to held-out clades

To assess transfer to clades absent from training at different taxonomic ranks, we constructed three PanTEon holdouts: the entire order Poales within kingdom Plantae, the entire phylum Arthropoda within kingdom Animalia, and the entire kingdom Fungi. The three holdouts therefore represent progressively greater phylogenetic distance from the training distribution. Clades were chosen *a priori* to span three ranks while leaving a trainable remainder with adequate superfamily coverage at each; no holdout was selected, substituted, or revised after any model was fit. For each holdout, GLARE, NeuralTE, and Terrier models were trained on the same partitions and evaluated on the same held-out sequences. Because clade-level exclusion removes near-identical family members shared with the training set, these holdouts also limit the effect of the record-level redundancy described in Section 2.1.1.

#### 2.2.3 Robustness to synthetic sequence degradation

To characterize robustness to the truncated and degraded TE copies that are common in genomes and in the *de novo* libraries built from them, we applied four sequence-degradation operators to the PanTEon benchmark test set, with training partitions left unmutated, and re-evaluated three classifiers on the mutated sequences. These operators impose controlled perturbations rather than calibrated models of TE decay. For GLARE, we reused the model from Section 2.2.1, evaluated in PanTEon’s native 30-superfamily, 8-order vocabulary, and NeuralTE and Terrier were trained through the PanTEon benchmark wrappers on the same partitions. All three classifiers were applied to the same mutated FASTA files derived from the 23,043 held-out test records, so every method scored identical sequences.

The four operators target different types of sequence damage. *Truncation* removes sequence so that the retained fraction is drawn as *f* ∼ Beta(*α, β*) with *α* = *µκ, β* = (1 ™ *µ*)*κ* and concentration *κ* = 10, so that each severity step yields a distribution of retained fractions rather than a fixed value, reproducing the heterogeneity expected among copies of differing age. The lost segment comes from one end, both ends, or the interior with probability 0.50*/*0.25*/*0.25. *Terminal damage* applies dense substitutions and indels within a window spanning a fraction *w* of each end, with intensity decaying as *e*^*™λd/wL*^ for distance *d* from the terminus, decay constant *λ* = 2.0, and element length *L*; the window is applied to the left, right, or both ends with probability 0.375*/*0.375*/*0.250. *Substitution* introduces per-position base substitutions at rate *s* (transition/transversion ratio 2.0; CpG sites mutated at 3× the base rate). *Indels* introduce per-position insertion and deletion events at rate *ι* (50% insertions; geometric length with mean 2 and maximum 10). All operator parameters were fixed *a priori*.

Two complementary designs were run. In the *isolated* design, each operator was swept alone across a five-step severity ladder while the other three were inactive, so that sensitivity to each type of damage could be measured separately. In the *combined* design, all four operators were applied together and increased in lockstep over five steps from Low to High, approximating the compounded decay of ancient elements. Ladder magnitudes spanned similar ranges of nominal severity, although severities are not directly comparable across operators: *µ* ran 0.90–0.50; *w* spanned 0.10–0.50 of each end, with within-window substitution 0.25–0.90 and indel event rate 0.015–0.08; *s* ran 0.04–0.20; and *ι* was tied to *s* at a fixed 1:5 ratio (0.008–0.040). An unmutated *intact* condition provided the shared baseline. Each mutated condition was generated in three independent technical replicates (distinct random seeds) and the intact baseline was evaluated once. Because element identity below the order level is not expected to survive heavy degradation, we report weighted order F1 in the main text and the corresponding weighted superfamily F1 in the Supplement; order labels were obtained from PanTEon’s superfamily-to-order map.

#### 2.2.4 Training database bias and cross-database transfer

To test whether performance estimates depend on training and evaluating within the same reference database, we separately fine-tuned three models on PanTEon only, Repbase only, and their union. Each was then evaluated on held-out PanTEon and Repbase records, yielding six train–test combinations. Order Poales was excluded from all three training sets and used as the evaluation clade for both databases. Poales is a well-defined partition in each, and holding it out means every model is tested on a clade it has not seen, so the comparison measures database transfer under generalization rather than within-distribution recall. To make predictions directly comparable, the label space of all models was restricted to the 25 superfamilies represented in both training sources.

### 2.3 Comparison of genomic language-model backbones

To identify a suitable backbone, we compared three pretrained genomic language models within a common GLARE classification and evaluation framework: DNABERT-2 [Zhou et al., 2023], Caduceus [Schiff et al., 2024], and NTv3 [Boshar et al., 2025]. This experiment used PanTEon’s native training/validation/test split shared across the three runs. All backbones were fully fine-tuned with a 512-nucleotide context, batch size 16, gradient accumulation over four minibatches, at most ten epochs, patience 8, label smoothing 0.05, backbone and head learning rates of 5 × 10^*™*5^ and 3 × 10^*™*4^, respectively, weight decay 0.01, and 5% warmup. These are conventional fine-tuning settings and were not tuned for any individual backbone, so differences in performance can be attributed to the backbones themselves. Using the same context in every run avoids confounding backbone identity with context length, although the window is shorter than the one used for the final model. Each classifier was optimized exclusively using superfamily cross-entropy, and validation and test sequences used 50%-overlapping window aggregation when longer than the context. Backbone selection was informed by the weighted superfamily F1 on held-out PanTEon sequences.

## 3 Results

### 3.1 GLARE classifies TEs accurately on held-out PanTEon and Repbase records

The final 32-superfamily GLARE model, trained with reverse-complement (RC) augmentation on the combined PanTEon and Repbase training data, was evaluated separately on the two source-specific test partitions (Table 2). Weighted superfamily F1 was 0.965 for the 23,043 PanTEon records and 0.946 for the 11,778 Repbase records. Weighted order F1 was 0.992 and 0.991, respectively, indicating that most superfamily-level errors remained within the correct order. Weighted by record count, the mean superfamily F1 across both sources was 0.958.

**Table 2:**
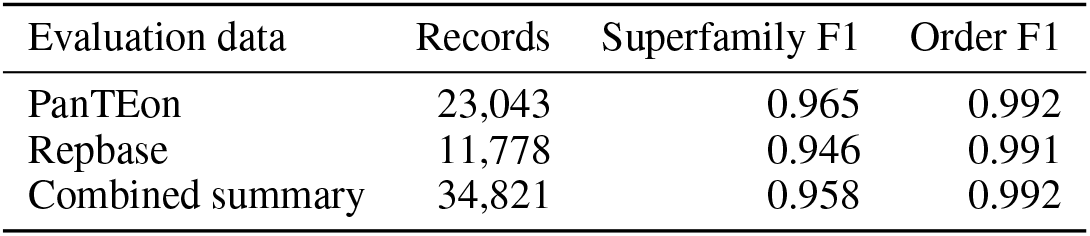
Final GLARE performance on held-out test records. Values are weighted F1-scores. The combined row is a record-count-weighted average of the two source-level summaries.

Performance was also largely independent of sequence orientation. In a matched ablation, the performance of an otherwise identical model trained without RC augmentation decreased by 0.047 in terms of weighted superfamily F1 when the same held-out records were reverse-complemented. In contrast, the score of the augmented model changed by only 0.001 (Supplementary Sections S1.2 and S2.1).

The combined PanTEon–Repbase confusion matrix was strongly diagonal (Figure 2), with the most frequent errors occurring between superfamilies within the same order. Among LTR elements, `LTR/COPIA` was predicted as `LTR/GYPSY` for 74 records, `LTR/GYPSY` was predicted as `LTR/COPIA` for 60 records, and 65 `LTR/OTHER` records were predicted as `LTR/GYPSY`. Among TIR elements, the most frequent off-diagonal assignment was `TIR/OTHER` to `TIR/HAT` (51 records). The heterogeneous `OTHER` categories had lower recall than many specific superfamilies: pooled recall was 0.757 for both `LTR/OTHER` (547/723) and `TIR/OTHER` (719/950), and 0.696 for `SINE/OTHER` (117/168), with 40 of these records assigned to SINE/TRNA. Some lower recall values were based on few examples, including `STRUCTURAL_RNA/STRUCTURAL_RNA` (6/9) and `TIR/P` (49/69), and are therefore uncertain.

**Figure 2.**
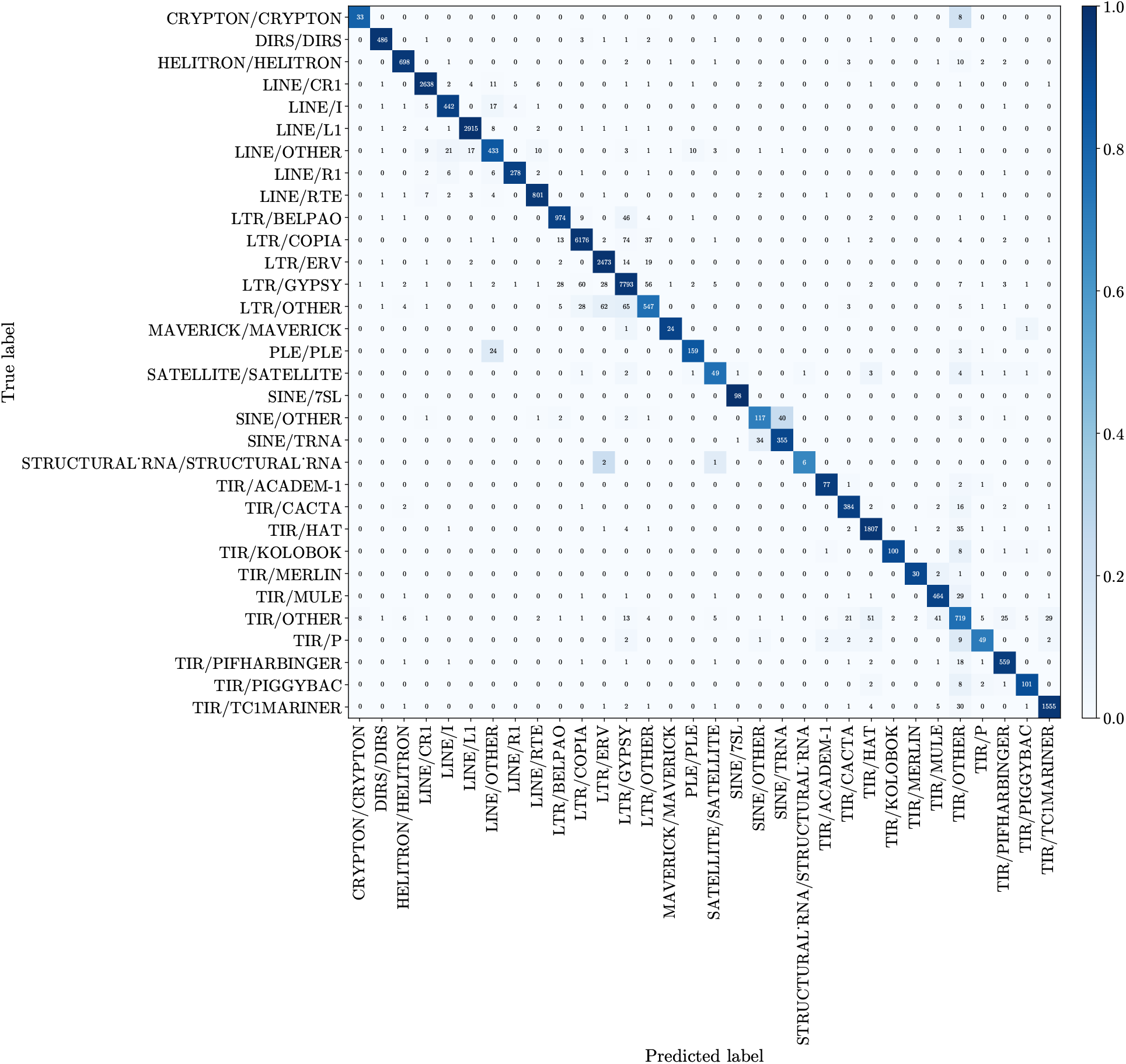
GLARE superfamily confusion matrix. Confusion matrix for the final 32-superfamily GLARE model on the combined held-out PanTEon and Repbase test records (*n* = 34,821). Rows indicate true labels and columns indicate predicted labels. Within each row, cell color represents the fraction of records assigned to each predicted label, whereas the overlaid annotations give pooled raw record counts. The source-specific matrices were aligned by superfamily label and summed cell by cell before row normalization.

Overall, the final model achieved high accuracy on both source-specific test sets, with residual errors concentrated among superfamilies within the same order and among heterogeneous OTHER categories. RC augmentation additionally removed nearly all of the orientation sensitivity observed without augmentation.

### 3.2 GLARE outperforms all nine classifiers on the PanTEon benchmark

For direct comparison with existing TE classifiers, GLARE was trained and evaluated under the PanTEon benchmark conditions, using PanTEon’s native 30-superfamily label space and the same record partitions used for the other methods. GLARE achieved a weighted superfamily F1 of 0.966 on the PanTEon test set (Table 3), the highest among the ten classifiers evaluated. The values reported by PanTEon for the nine retrained TE classifiers ranged from 0.539 for BERTE to 0.922 for NeuralTE, giving GLARE an absolute margin of 0.044 over the next-highest method.

**Table 3:**
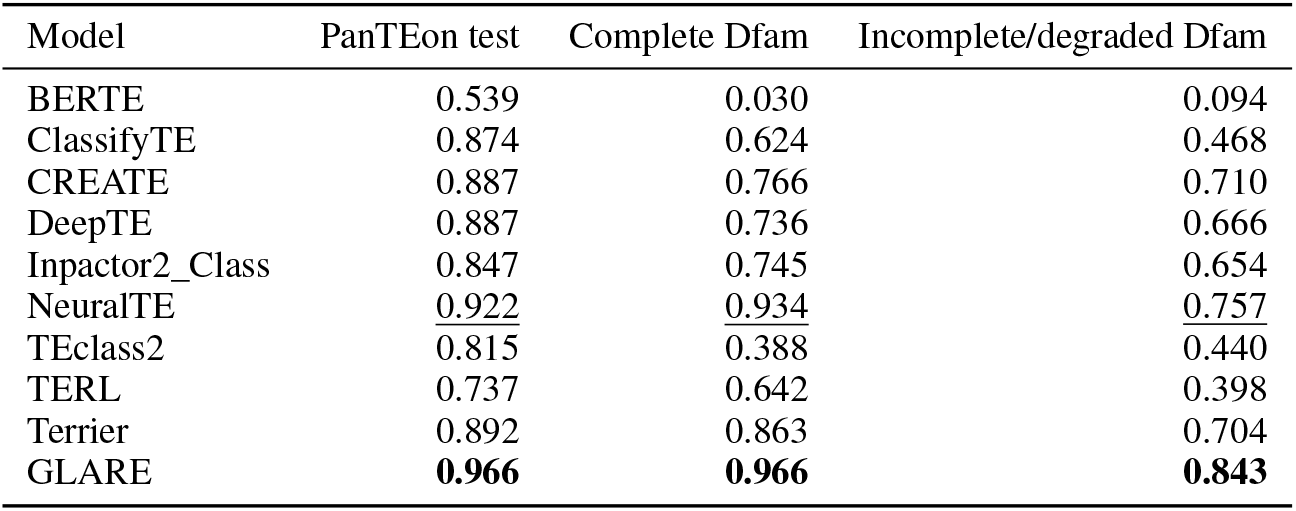
Weighted superfamily F1 across the PanTEon benchmark evaluations. Results are shown for GLARE and all nine classifiers included in the PanTEon benchmark. The Dfam evaluation sets contain complete and incomplete or degraded elements from species absent from the PanTEon training data. Boldface indicates the highest value in each column, and underlining indicates the second highest.

This advantage was retained on the two unseen-species Dfam evaluation sets. On the 867 complete elements, GLARE achieved a weighted superfamily F1 of 0.966, compared with 0.934 for NeuralTE, the next-highest classifier. On the 1,442 incomplete or degraded elements, GLARE achieved 0.843, compared with 0.757 for NeuralTE. Performance was lower on the incomplete or degraded elements than on the complete elements for GLARE and most comparison methods, but GLARE retained the highest F1 on both datasets.

Thus, GLARE achieved the highest weighted superfamily F1 on the PanTEon test set and both independently curated Dfam datasets, including elements from species absent from the PanTEon training data.

### 3.3 GLARE retains the highest accuracy on held-out clades

To evaluate taxonomic transfer, GLARE, NeuralTE, and Terrier were retrained separately for three PanTEon clade holdouts using the same clade-specific partitions. The holdouts spanned three taxonomic ranks: Poales excluded an entire order within Plantae, Arthropoda an entire phylum within Animalia, and Fungi an entire kingdom. GLARE achieved the highest weighted superfamily F1 point estimate in all three evaluations (Figure 3), scoring 0.897 for Poales, 0.887 for Arthropoda, and 0.783 for Fungi. This ordering is consistent with increasing phylogenetic distance from the training distribution across the three evaluated holdouts.

**Figure 3.**
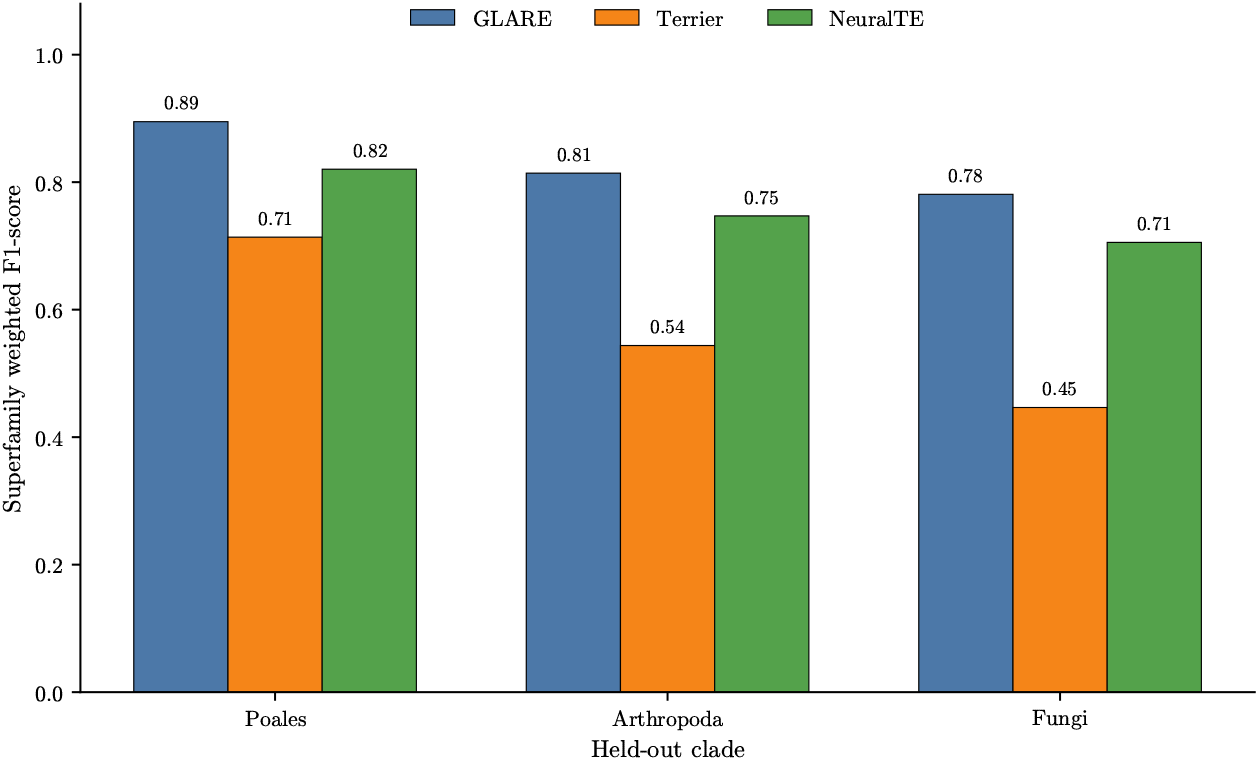
Transposable element classification in unseen clades. Bars report weighted superfamily F1 for GLARE, NeuralTE, and Terrier after separately excluding all records from the order Poales, the phylum Arthropoda, or the kingdom Fungi from the corresponding training set. For each holdout, all three classifiers were retrained and evaluated using the same clade-specific partitions.

For the Poales order holdout, NeuralTE and Terrier obtained 0.820 and 0.714, respectively. Their corresponding scores dropped to 0.747 and 0.544 for the Arthropoda phylum holdout, and to 0.706 and 0.447 for the Fungi kingdom holdout. Like GLARE, both comparison methods obtained their lowest score on the Fungi holdout, with a larger decrease for Terrier. Relative to NeuralTE, GLARE’s absolute F1 margins were 0.077 for Poales, 0.14 for Arthropoda, and 0.077 for Fungi; the corresponding margins over Terrier were substantially larger at 0.183, 0.343, and 0.336.

Thus, although all three classifiers declined across the increasingly distant clade holdouts, GLARE retained the highest weighted superfamily F1 in every evaluation. Its advantage over both comparison methods persisted even when the entire kingdom Fungi was excluded from training.

### 3.4 GLARE is the most robust classifier under synthetic sequence degradation

To evaluate robustness to sequence decay, GLARE, NeuralTE, and Terrier were applied to identical degraded versions of the PanTEon benchmark test records. Because severe degradation may obscure superfamily identity while preserving the broader TE order, weighted order F1 was used as the primary metric. When all four degradation operators were applied together, every classifier declined along the severity ladder, but GLARE retained the highest weighted order F1 at every step (Figure 4). From an intact baseline of 0.992, GLARE declined to 0.977 at Low, 0.838 at Medium, and 0.494 at High severity. NeuralTE and Terrier declined more steeply from their respective intact baselines, reaching 0.475 and 0.347 at High severity. GLARE’s advantage over NeuralTE, the next-highest classifier, was greatest at intermediate severity, where it reached 0.104, before narrowing to 0.019 at High severity as performance declined for all three methods.

**Figure 4.**
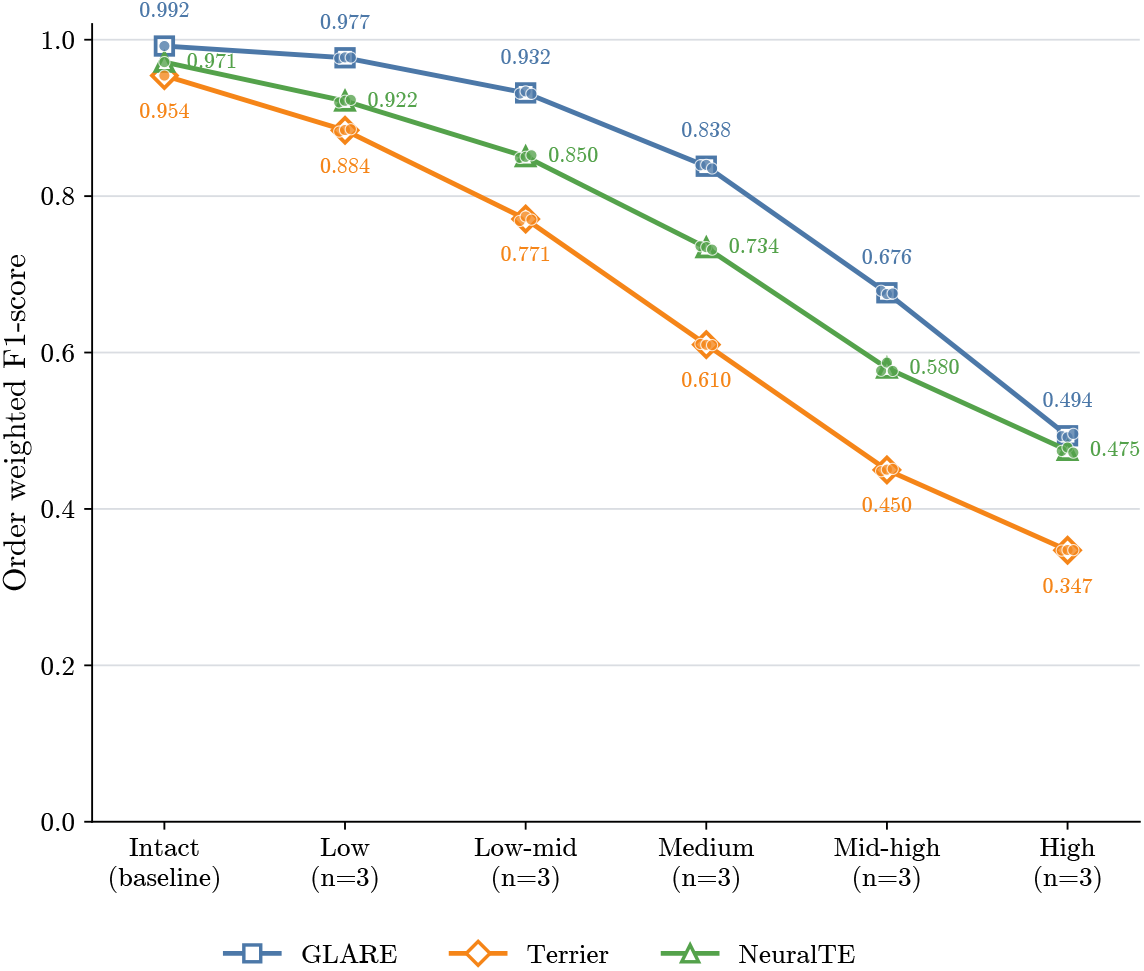
Classifier robustness to compounded synthetic degradation. Weighted order F1 of GLARE, NeuralTE, and Terrier on the PanTEon benchmark test set with all four degradation operators applied together at increasing severity (Section 2.2.3). Lines show the mean of three technical replicates; small points show individual replicates. Superfamily-level results and the isolated single-operator evaluations are shown in Supplementary Figures S2–S4.

The corresponding weighted superfamily F1 declined more steeply for every method (GLARE 0.966 intact to 0.230 at High severity; Supplementary Figure S2). GLARE’s confusion matrices showed that this divergence between order- and superfamily-level performance was driven largely by errors within the LTR order. In PanTEon’s native vocabulary, LARD (Large Retrotransposon Derivative) denotes a non-autonomous, internally deleted LTR class. As degradation increased, autonomous LTR elements were increasingly assigned to this class: GYPSY→LARD and COPIA→LARD were the dominant confusions at Medium severity, and this pattern persisted at higher severities (Supplementary Figure S5). Because all three classes belong to the LTR order, this collapse toward a degradation-like within-order class left order calls largely intact even as superfamily resolution was lost.

The isolated single-operator sweeps showed the same overall ranking: GLARE retained the highest weighted order F1 of the three classifiers under every operator and at every severity tested (Supplementary Figure S3). The same pattern held at the superfamily level (Supplementary Figure S4). Terrier was the most sensitive classifier to truncation, terminal damage, and substitution, with a particularly large decline under dense substitution (0.577 weighted superfamily F1 at a per-position rate of 0.20, compared with 0.842 for GLARE). Indels were the only operator under which NeuralTE (0.840) fell below Terrier (0.870) at the highest severity.

Thus, GLARE was the most robust of the three classifiers under both compounded and isolated degradation conditions. Its advantage across truncation, terminal damage, substitution, and indels indicates that the observed robustness was not restricted to a single form of sequence damage.

### 3.5 Cross-database transfer is asymmetric and combined training closes the gap

To test whether the training database affects performance, GLARE models were trained on PanTEon only, Repbase only, or both combined, and evaluated on held-out Poales records from each database. When trained and evaluated on records from the same source database, the PanTEon-only and Repbase-only models achieved similar weighted superfamily F1 values of 0.943 and 0.940, respectively (Figure 5). Cross-database transfer was nevertheless strongly asymmetric. The PanTEon-only model declined from 0.943 on held-out PanTEon Poales records to 0.813 on Repbase Poales records, a difference of 0.130. Conversely, the Repbase-only model scored 0.940 on Repbase and 0.927 on PanTEon, a difference of only 0.013. Similar within-database scores therefore did not imply similar transfer to the other database.

**Figure 5.**
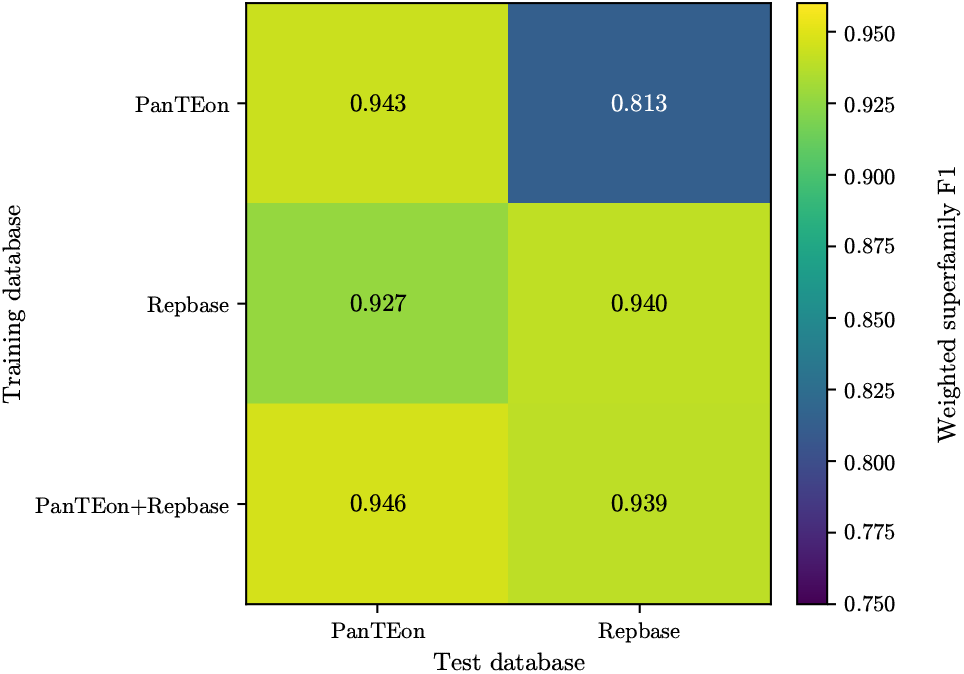
Cross-database weighted superfamily F1. Rows indicate training on PanTEon only, Repbase only, or combined PanTEon and Repbase data; columns indicate evaluation on held-out PanTEon or Repbase Poales records. All six evaluations used the shared 25-superfamily label space.

Training on the combined databases largely eliminated this imbalance. The combined model achieved the highest score on PanTEon (0.946) and a score of 0.939 on Repbase, only 0.001 below the Repbase-only model. Relative to PanTEon-only training, the addition of Repbase substantially improved performance on Repbase while also slightly improving performance on the PanTEon test set itself. Combined training thus led to near-best performance on both databases and removed most of the transfer loss seen with PanTEon-only training.

### 3.6 NTv3 backbone outperforms DNABERT-2 and Caduceus

To compare backbones without architecture-specific hyperparameter optimization, DNABERT-2, Caduceus, and NTv3 were fine-tuned using the same PanTEon split, 512-nucleotide context, and fixed training settings (Table 4). NTv3 achieved the highest weighted superfamily F1 (0.958) and order F1 (0.990) on the PanTEon test partition, followed by DNABERT-2 (0.928 and 0.973, respectively). Caduceus performed considerably worse, with a superfamily F1 of 0.817 and an order F1 of 0.920. Based on this comparison, NTv3 was selected as the backbone for GLARE.

**Table 4:**
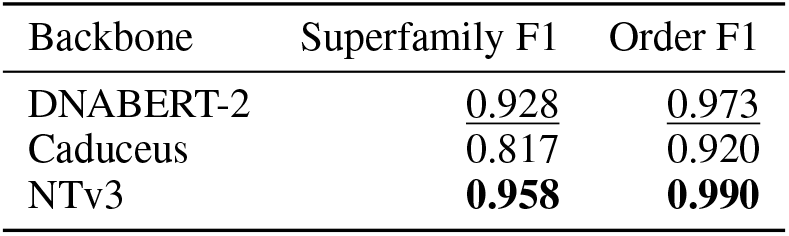
Fixed-setting comparison of genomic language-model backbones. All backbones used the same PanTEon data split, 512-nucleotide context, and training settings. Values are weighted F1-scores on the PanTEon test partition.

| Backbone | Superfamily F1 | Order F1 |
| --- | --- | --- |
| DNABERT-2 | 0.928 | 0.973 |
| Caduceus | 0.817 | 0.920 |
| NTv3 | **0.958** | **0.990** |

## 4 Discussion

GLARE fine-tunes a genomic language model on harmonized PanTEon and Repbase reference data to classify transposable elements from sequence alone. In the standardized PanTEon benchmark, GLARE achieved the highest superfamily F1 among all classifiers tested, and its advantage over other tools was retained when its generalization was tested on unseen taxonomic clades and degraded TE sequences. The contributions here are both conceptual and practical. Conceptually, this study is, to our knowledge, the first demonstration of a fine-tuned genomic language model applied to TE classification. Practically, GLARE is an easy-to-use, stand-alone classifier that assigns labels directly to candidates in a *de novo* repeat library without requiring a close reference match or a prescribed set of diagnostic features.

Our primary evaluation used the recently published PanTEon benchmarking framework, which retrains existing classifiers on the same records and evaluates them with the same labels and metric [Orozco-Arias et al., 2026]. Before PanTEon, published TE classifiers differed in training data, label hierarchies, and data splits, so their reported scores were not directly comparable. PanTEon also tests generalization on manually curated Dfam libraries from species absent from its training data, divided into (a) complete and (b) incomplete or degraded elements. GLARE ranked first on all of these test sets, and its margin over the other tools was largest on the most challenging of them, the incomplete or degraded elements. This is particularly relevant to *de novo* repeat annotation, where degraded copies are common. The strongest alternative, NeuralTE, explicitly encodes target site duplications, terminal repeats, and protein domains [Hu et al., 2024]; GLARE outperformed it without being given any of these features. Our remaining evaluations extend PanTEon’s framework to further tests of generalization: the evolutionary distance between training and test sequences, the type and extent of sequence damage, and the database a model is trained on.

PanTEon’s Dfam evaluation excludes species seen in training; however, close relatives of training species can still be present in the test set. Withholding entire taxonomic clades at the order, phylum, and kingdom levels explicitly sets how distant the test sequences are from the training set. Beyond testing the models’ generalization, this has practical implications, as TE annotation is often done in studies of species from under-represented clades, with few or no TEs from close relatives in reference libraries. GLARE outperformed NeuralTE and Terrier in the Poales, Arthropoda, and Fungi holdouts, suggesting that its learned representation retains superfamily signal that transfers across large evolutionary distances.

PanTEon’s evaluation on incomplete or degraded Dfam elements and our synthetic-mutation experiments examine generalization to degraded elements in complementary ways. The Dfam evaluation uses real degraded elements but does not control the level or type of degradation, which our synthetic test varies systematically. GLARE was the most accurate classifier in both, ranking first in the synthetic test under every degradation operator and at every severity. As damage increased, the gap between order and superfamily accuracy widened. Although GLARE was trained only on superfamily labels, its superfamily errors largely stayed within the correct order. This suggests that the model learned order-level similarity from sequence alone. The large number of GYPSY and COPIA elements predicted as LARD (large retrotransposon derivatives) is particularly interesting here. Because LARDs evolve from autonomous LTR retrotransposons through the loss of their internal coding regions [Kalendar et al., 2004], the synthetic mutation of LTR elements may partially simulate this process, making these misclassifications consistent with how LARDs arise.

Reference databases differ in which species they cover, how they curate elements, and how they label them, so a classifier trained on one may not transfer well to another. To quantify this effect, we tested how well models trained on Repbase performed on PanTEon records, and vice versa. As in the Poales holdout above, order Poales was withheld from training, so database transfer was measured under generalization to an unseen clade. Models trained on PanTEon or Repbase scored similarly on records from their own source, but the PanTEon-only model transferred less well to Repbase than the Repbase-only model did to PanTEon. One possible explanation is that Repbase, which has undergone years of manual curation, contains a more diverse and challenging set of TEs. Nevertheless, the PanTEon-only model did not collapse on Repbase. Training the final model on both resources was a practical choice to make it as robust as possible. The combined model scored similarly to the single-source models on their own databases. Adding a second source therefore extended coverage to both databases without sacrificing accuracy on either.

All classifiers in the PanTEon benchmark were trained on the same partitions, but only GLARE also underwent a hyperparameter search on the validation set. Although the test set was never used in this search, validation and test records come from the same corpus, so part of GLARE’s lead could in principle reflect this extra optimization step rather than the model itself. GLARE’s performance on the clade holdouts and the Dfam sets already argues against this, but the fixed-setting backbone comparison, run to choose a genomic language model for GLARE, addresses the concern more directly. Even before hyperparameter optimization, using the GLARE architecture with generic fine-tuning settings, an untuned classification head, and a 512-nucleotide context, two of the three fine-tuned backbones exceeded every previously reported PanTEon score. NTv3 reached 0.958 and DNABERT-2 0.928, compared with 0.922 for NeuralTE. Hyperparameter optimization and the longer context together raised NTv3’s score by only 0.008, to 0.966. GLARE’s lead on the PanTEon benchmark therefore does not depend on hyperparameter tuning but comes mainly from fine-tuning a suitable pretrained backbone. The model with Caduceus as the backbone reached only 0.817 under the same settings, however, so the choice of the underlying genomic language model still matters.

This study has several limitations. The use of record-level random splits in the PanTEon benchmark can place closely related sequences in both the training and the test sets. The clade holdouts avoid this by withholding entire taxonomic clades from training, but only one clade was tested at each taxonomic rank. The synthetic degradation modes were chosen *a priori* rather than fitted to mutation rates and patterns estimated from naturally degraded copies. The backbone comparison reveals that the choice of genomic language model matters. However, because the backbones differ in architecture, size, tokenization, and pretraining data, the comparison does not show which of these makes NTv3 the stronger choice. As a classification tool, GLARE does not detect TEs or refine their boundaries. Its performance on *de novo* repeat libraries generated within complete annotation workflows remains to be tested; such repeat libraries contain elements longer than the model context, sequences in either orientation, and many sequences that are not TEs at all. GLARE handles the first two through overlapping windows and reverse-complement augmentation, and the third through satellite and structural RNA classes and a confidence threshold that labels uncertain calls as UNKNOWN. However, this threshold is not a validated out-of-distribution test; that would require training with explicit unknown classes and choosing the threshold on held-out data that contains them.

GLARE’s advantage over other classifiers does not depend on hyperparameter tuning, so newer backbones can be fine-tuned in the same way. Other reference databases can be added through label mapping, as we did with Repbase, which removed most of the cross-database loss without lowering accuracy on PanTEon. GLARE can therefore be readily updated as genomic language models and TE reference libraries improve.

## 5 Supplementary Materials

Supplementary Section S1.1 and Table S1 report the hyperparameter-optimization search domain and the selected value for each dimension. Supplementary Sections S1.2 and S2.1, together with Supplementary Table S2 and Supplementary Figure S1, report the reverse-complement augmentation ablation. Supplementary Section S2.2 and Supplementary Figures S2–S5 report the superfamily-level combined ladder, the isolated single-operator sweeps, and GLARE’s superfamily confusion matrices along the combined ladder for the synthetic-degradation experiment. Supplementary Files S1–S2 give the source-specific mappings from PanTEon and Repbase categories to the GLARE label space.

## 6 Data Availability

GLARE is available at code.jgi.doe.gov/NHaas/glare.

## 7 Acknowledgements

The work conducted by the U.S. Department of Energy Joint Genome Institute (https://ror.org/04xm1d337), a DOE Office of Science User Facility, is supported by the Office of Science of the U.S. Department of Energy operated under contract no. DE-AC02-05CH11231. Supported by the Laboratory Directed Research and Development (LDRD) program at Lawrence Berkeley National Laboratory.

