## Supplemental Material for "GLARE: Genomic Language model for Automated Repetitive Element classification"

Supplementary Materials

Nicolas Haas*[1] and Tomáš Brůna†[1]
[1]DOE Joint Genome Institute, Lawrence Berkeley National Laboratory,
1 Cyclotron Road, Berkeley, CA 94720, USA

October 6, 2026

### S1 Supplementary Methods

#### S1.1 Hyperparameter search domain

Table S1 reports the search domain and the selected value for each dimension of the hyperparameter optimization described in the main text.

Table S1: Hyperparameter-optimization (HPO) search dimensions and values selected for the final GLARE model.

| Hyperparameter | Search domain | Selected value |
|---|---|---|
| Backbone learning rate | $[10^{-6}, 10^{-4}]$ | $2.5303 \times 10^{-5}$ |
| Head learning rate | $[5 \times 10^{-5}, 5 \times 10^{-4}]$ | $2.8944 \times 10^{-4}$ |
| Weight decay | $[0, 0.1]$ | 0.05970 |
| Batch size | {4, 8} | 8 |
| Label smoothing | $[0, 0.15]$ | 0.04655 |
| Class-weighting scheme | None; Square-root inverse frequency; Inverse frequency | Square-root inverse frequency |
| Maximum sequence length | {512, 1,024, 2,048, 4,096, 8,192} | 4,096 |
| Sliding-window overlap | {0, 0.25, 0.5} | 0.5 |
| Hidden dimension | {256, 512, 1,024} | 512 |
| Head dropout | $[0, 0.3]$ | 0.2232 |

#### S1.2 Ablation of reverse-complement augmentation

Strand-orientation robustness was evaluated in the combined PanTEon–Repbase, 32-superfamily setting without a taxonomic holdout. Two otherwise matched NTv3 models were trained with reverse-complement (RC) augmentation disabled or enabled. When enabled, each training example was reverse-complemented independently with probability 0.5, after any random long-sequence crop, while retaining its order and superfamily labels.

Each trained model was evaluated on the original orientation and on a deterministic RC-only transformation of the same held-out PanTEon and Repbase random-test records. The principal analysis was a $2 \times 2$ matrix crossing two

*
†

training exposures (original-only versus original+RC) with two test orientations (original versus RC-only). For each cell, PanTEon and Repbase confusion-matrix counts were aligned by label and pooled before weighted superfamily F1 was recomputed. RC-only evaluation reverse complemented every test sequence before tokenization and did not create a distinct random sample.

### S2 Supplementary Results

#### S2.1 Reverse-complement augmentation reduces orientation sensitivity

Without RC augmentation, reversing all held-out sequences reduced weighted superfamily F1 from 0.965 to 0.938 on PanTEon and from 0.945 to 0.862 on Repbase (Table S2; absolute changes of $-0.027$ and $-0.082$). With original+RC training, the same paired comparisons changed from 0.965 to 0.966 on PanTEon and from 0.946 to 0.948 on Repbase. RC augmentation therefore increased RC-only F1 by 0.028 on PanTEon and 0.085 on Repbase, while changing original-orientation F1 by less than 0.002 in either database.

Table S2: Weighted superfamily F1 across training exposure and test orientation. Original and RC-only tests contain the same held-out records in opposite orientations.

| Evaluation data | Training exposure | Original test | RC-only test | RC–Original |
|---|---|---|---|---|
| PanTEon | Original-only | 0.965 | 0.938 | −0.027 |
| PanTEon | Original+RC | 0.965 | 0.966 | +0.001 |
| Repbase | Original-only | 0.945 | 0.862 | −0.082 |
| Repbase | Original+RC | 0.946 | 0.948 | +0.001 |
| PanTEon+Repbase | Original-only | 0.958 | 0.911 | −0.047 |
| PanTEon+Repbase | Original+RC | 0.958 | 0.959 | +0.001 |

In the principal pooled $2 \times 2$ analysis (Figure S1), original-only training decreased from 0.958 on original-orientation records to 0.911 on their RC-only versions, whereas original+RC training yielded 0.958 and 0.959, respectively.

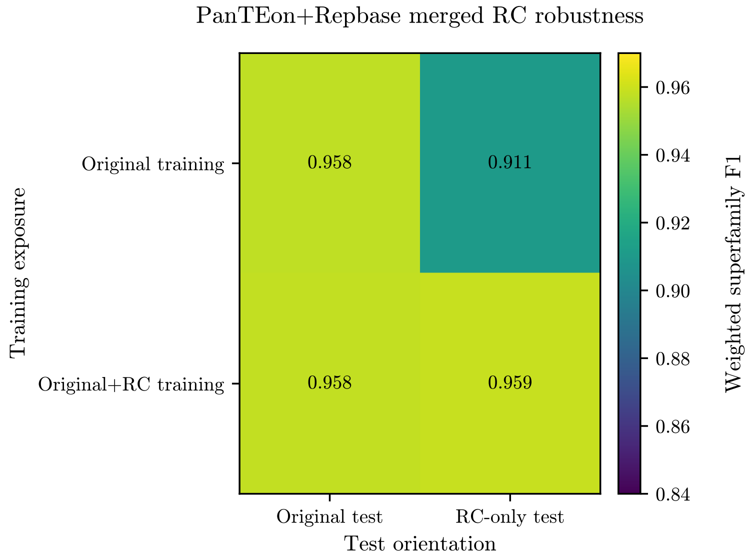


Figure S1: Principal $2 \times 2$ reverse-complement robustness analysis after pooling PanTEon and Repbase held-out counts. Rows cross original-only versus original+RC training exposure; columns cross original versus RC-only presentation of the same test records. Colors and annotations report weighted superfamily F1 recomputed from label-aligned pooled confusion-matrix counts.

#### S2.2 Robustness to synthetic sequence degradation

The synthetic-degradation experiment is described in the main-text Materials and Methods, and the main-text figure reports weighted order F1 for the combined all-four-operator ladder. Here we provide the corresponding weighted superfamily F1 for the same combined ladder (Figure S2), the isolated single-operator sweeps at both the order (Figure S3) and superfamily (Figure S4) levels, and GLARE's superfamily confusion matrices at three points along the combined ladder within the LTR order (Figure S5). Superfamily scores declined more steeply than order scores for every classifier and at every severity, consistent with degraded elements retaining order-level but not superfamily-level identity. GLARE nonetheless retained the highest weighted F1 at all levels, under the combined ladder and under each operator applied alone.

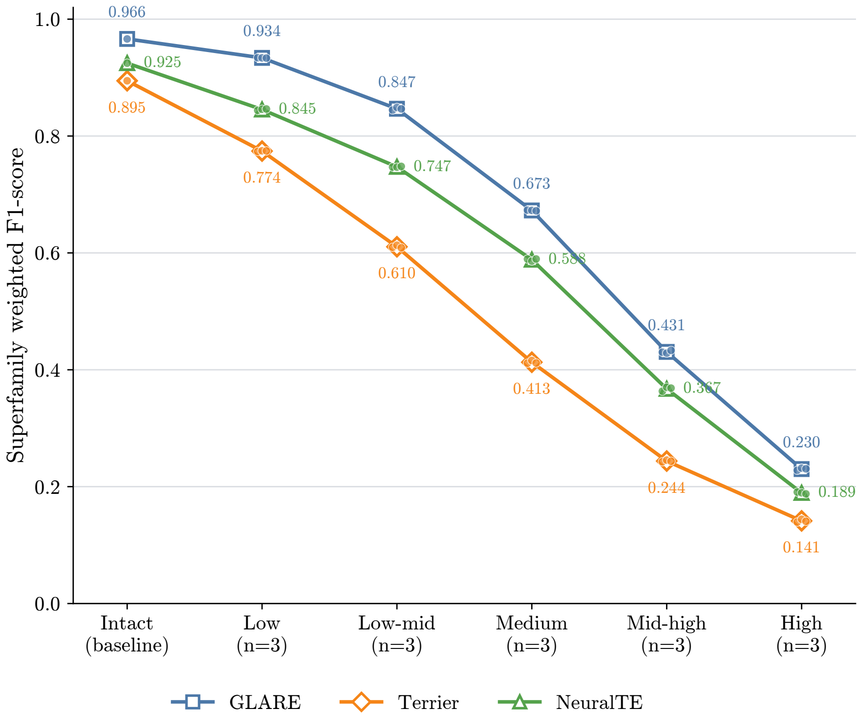


Figure S2: Weighted superfamily F1 under the combined four-operator degradation ladder, for GLARE, Terrier, and NeuralTE on the PanTEon benchmark test set. Companion to main-text Figure 4, which reports weighted order F1 for the same conditions. Lines and large markers are means over three technical replicates per mutated rung (one shared intact baseline); small points are the individual replicates.

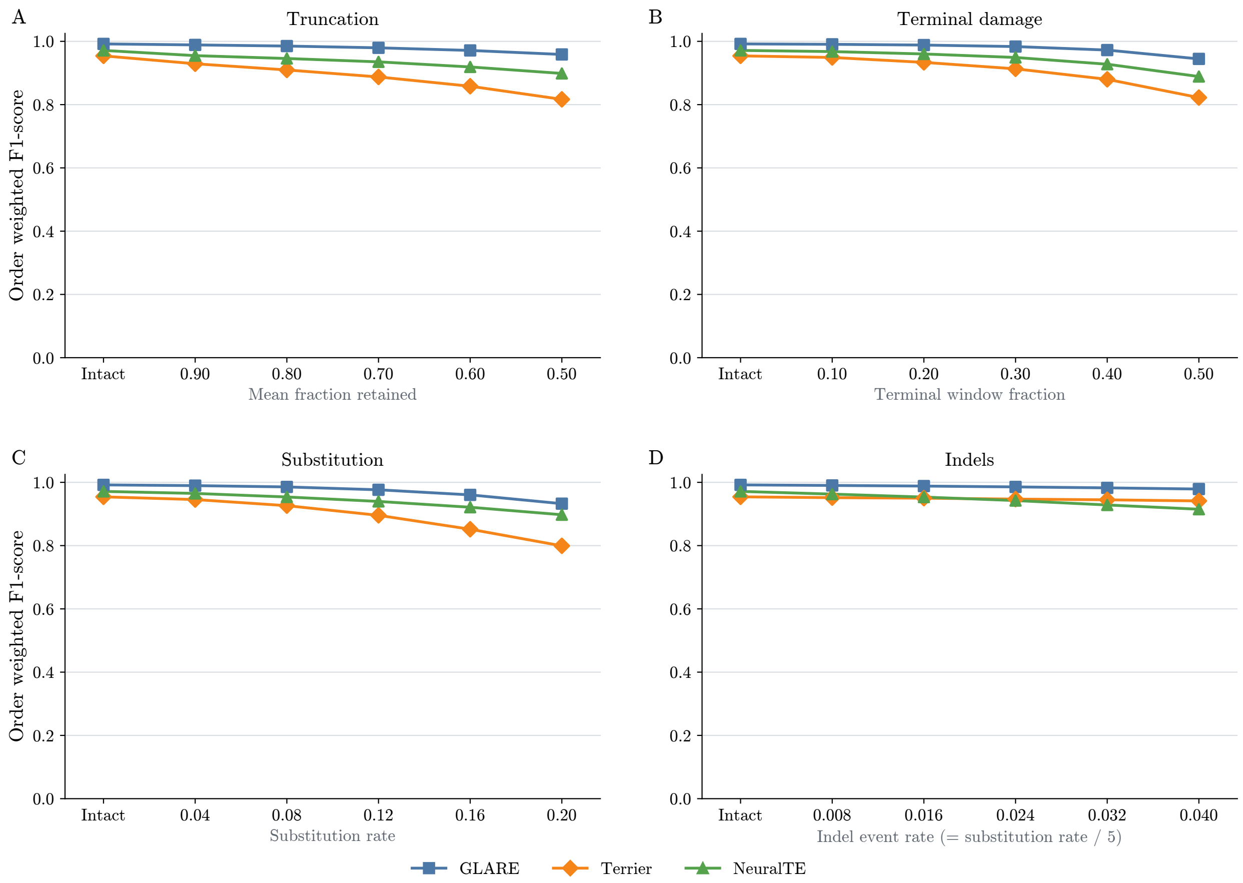


Figure S3: Weighted order F1 when each degradation operator is swept alone while the others are inactive, on the PanTEon benchmark test set. (A) Truncation (mean fraction retained 0.90–0.50). (B) Terminal damage (window fraction 0.10–0.50; within-window substitution 0.25–0.90, indel event rate 0.015–0.08). (C) Substitution (per-position rate 0.04–0.20; ti/tv 2.0, CpG 3×). (D) Indels (per-position event rate 0.008–0.040, tied to the substitution rate at 1:5). Operator definitions are given in the main-text Materials and Methods. The leftmost point in each panel is the intact baseline; markers are means over three technical replicates.

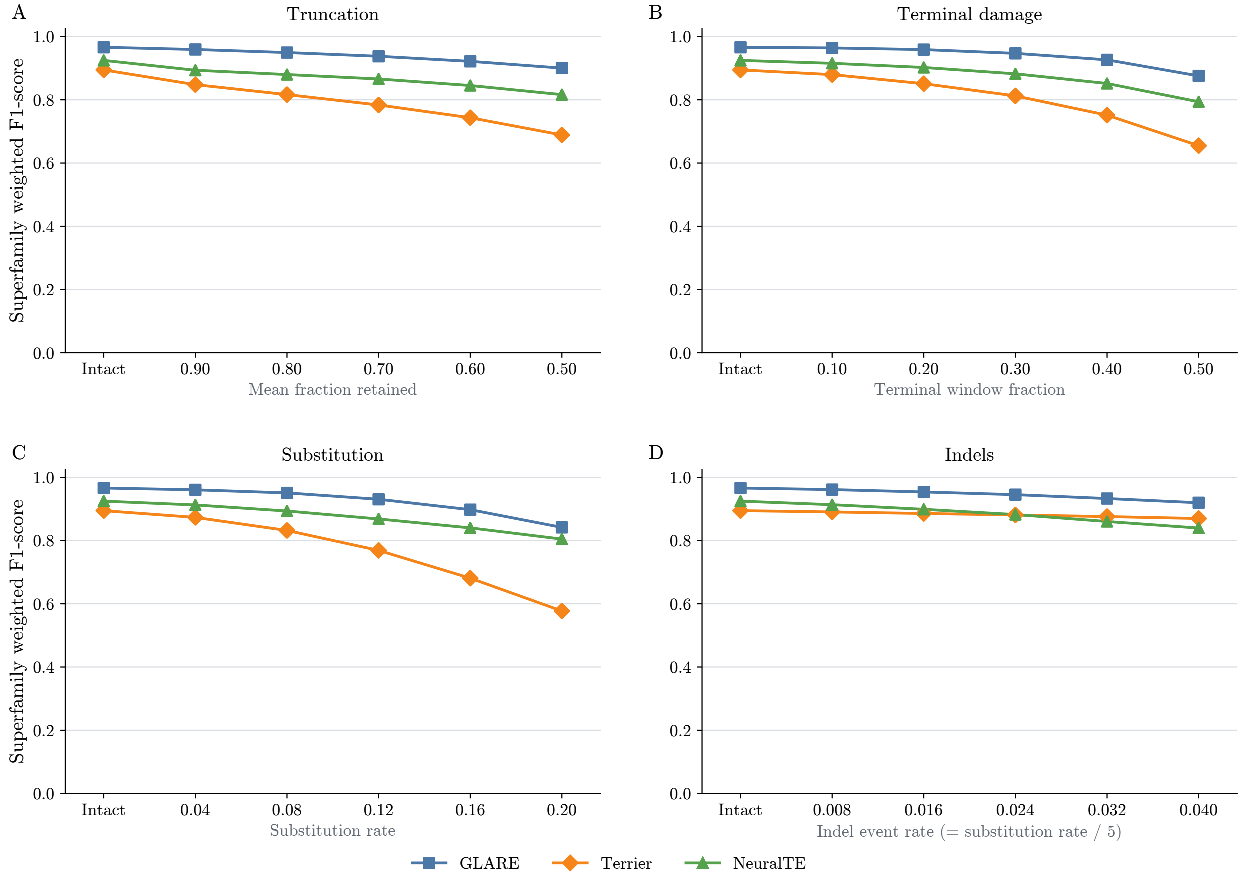


Figure S4: Weighted superfamily F1 for the isolated single-operator sweeps, with panels, operators, and magnitudes as in Figure S3.

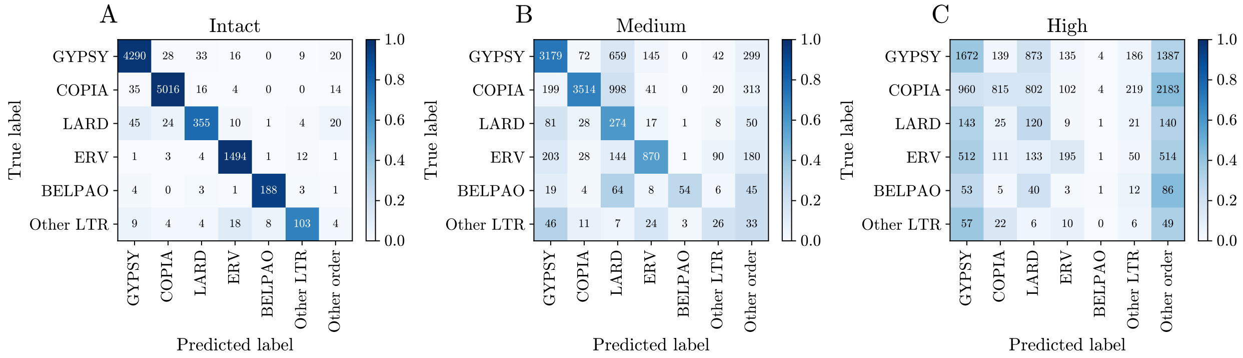


Figure S5: GLARE superfamily confusion matrices for LTR elements under the combined degradation ladder, at the intact baseline (A), Medium severity (B), and High severity (C), on the PanTEon benchmark test set. Rows are the true LTR superfamilies; columns are the predicted LTR superfamilies plus a single `Other order` column collecting all predictions outside the LTR order. Cell color is normalized within each true-label row and therefore represents recall allocation, whereas annotations are raw record counts. Records whose true label lies outside the LTR order are not shown. Mutated panels show one representative replicate. Mass moves off the diagonal into the `LARD` column at Medium severity while the `Other order` column remains sparse; at High severity the `Other order` column is also populated, reflecting the loss of order-level accuracy at the harshest rung
